# A Bayesian Method to Incorporate Hundreds of Functional Characteristics with Association Evidence to Improve Variant Prioritization

**DOI:** 10.1101/000984

**Authors:** Sarah A. Gagliano, Michael R. Barnes, Michael E. Weale, Jo Knight

## Abstract

The increasing quantity and quality of functional genomic information motivate the assessment and integration of these data with association data, including data originating from genome-wide association studies (GWAS). We used previously described GWAS signals (“hits”) to train a regularized logistic model in order to predict SNP causality on the basis of a large multivariate functional dataset. We show how this model can be used to derive Bayes factors for integrating functional and association data into a combined Bayesian analysis. Functional characteristics were obtained from the Encyclopedia of DNA Elements (ENCODE), from published expression quantitative trait loci (eQTL), and from other sources of genome-wide characteristics. We trained the model using all GWAS signals combined, and also using phenotype specific signals for autoimmune, brain-related, cancer, and cardiovascular disorders. The non-phenotype specific and the autoimmune GWAS signals gave the most reliable results. We found SNPs with higher probabilities of causality from functional characteristics showed an enrichment of more significant p-values compared to all GWAS SNPs in three large GWAS studies of complex traits. We investigated the ability of our Bayesian method to improve the identification of true causal signals in a psoriasis GWAS dataset and found that combining functional data with association data improves the ability to prioritise novel hits. We used the predictions from the penalized logistic regression model to calculate Bayes factors relating to functional characteristics and supply these online alongside resources to integrate these data with association data.

**Author Summary:** Large-scale genetic studies have had success identifying genes that play a role in complex traits. Advanced statistical procedures suggest that there are still genetic variants to be discovered, but these variants are difficult to detect. Incorporating biological information that affect the amount of protein or other product produced can be used to prioritise the genetic variants in order to identify which are likely to be causal. The method proposed here uses such biological characteristics to predict which genetic variants are most likely to be causal for complex traits.

## Introduction

Genome-wide association studies (GWAS), which investigate the association between genetic variation and phenotypic traits, have identified many genes associated with human diseases [1]. However, despite considerable advances, much of the estimated heritability remains unaccounted for. Purcell et al. [2] showed that single nucleotide polymorphisms (SNPs) from GWAS with sub-threshold p-values account for a considerable proportion of the variance in independent samples suggesting that they are enriched for causal SNPs or their proxies. The issues of small sample size, low minor allele frequency, and lack of linkage disequilibrium (LD) between genotyped SNPs and the un-genotyped causal SNPs present challenges to detecting truly causal variants among near-significant genetic associations.

Emerging experimental data from various sources have suggested that the functional characteristics of specific genomic regions, such as histone modifications, DNase I hypersensitive sites, transcription factor binding sites, and expression quantitative trait loci (eQTL) among others, could offer biological explanations for many variants found to be associated with disease [3, 4, 5, etc.]. In September 2012, a series of publications from the ENcyclopedia of DNA Elements (ENCODE) Project Consortium, had the key message that approximately 80% of the human genome, including non-coding and intergenic regions, overlaps with at least one functional element that may be active in certain cell types, under defined physiological conditions [6]. Furthermore, putative disease-causing variants show significant enrichment for multiple functional characteristics from the ENCODE Project [7]. For example, GWAS variants or variants with which they are in perfect LD are more frequently localized to DNase I hypersensitive sites than would be expected by chance [8].

Various tools are available that allow one to summarise the functional characteristics of variants in a given region. For instance, Boyle et al. developed RegulomeDB, a web-based interface that provides an easily interpretable score from an amalgamation of many functional characteristics derived from a variety of sources to annotate non-coding variants [9]. Other programs such as HaploReg [10] and SNPnexus [11] perform similar functions and account for LD. Although these programs provide facile access to summary information about the location of variants, they are only able to provide a relatively arbitrary and crude ranking of functional significance. The ranking scale used in RegulomeDB is based on the number of categories into which a variant falls with the highest scores given to those variants that fall into both an eQTL and a transcription factor binding site, regardless of cell type or specific transcription factor.

The central challenge in the interpretation of genetic associations lies in the processing and meaningful integration of a hugely diverse range of information. Having derived a score for a region containing a candidate variant, it has to be integrated with association evidence. We proposed the use of empirically derived weightings within a Bayesian framework [3]. More recently Schork et al. suggested the use of stratified False Discovery Rate (sFDR) and Darnell et al. proposed multi-thresholding in a manner that they say is equivalent to varying the significance threshold at each marker depending on the prior information [12,13]. In order to implement these approaches one needs to define appropriate weights. For instance, Schork et al. [12] used an LD-weighted scoring algorithm, and Kindt et al. [14] recently published a multivariate logistic regression approach. However, neither of these approaches is easily scalable to the very large number of functional characteristics that are becoming available.

The primary objectives of this study are to describe an empirically justified method to identify which functional characteristics are best correlated with GWAS hit SNPs, to provide clues to the etiology of such traits, and to develop and implement a method to incorporate functional characteristics with statistical information in association studies. To achieve these objectives we use a machine learning approach, elastic net (a regularized logistic regression), to predict causality of a SNP based on information from 543 functional characteristics. We explore models based on all GWAS significant SNPs and also subsets of significant SNPs selected on the basis of phenotype and p-value. Functional characteristics are considered individually or in groups. We report a) the accuracy of the predictions to demonstrate the utility of the method and to investigate the behaviour of the different models, b) the frequency, correlation between and coefficients of the functional characteristics providing insight about their functional relevance to disease, c) a prediction score for each SNP, and d) details of how to combine this score with association statistics in a formal Bayesian framework.

We provide online scripts that can be employed so the method can be used by other researchers using additional functional characteristics (http://www.camh.ca/en/research/research_areas/genetics_and_epigenetics/Pages/Statistical-Genetics.aspx). For the best models we provide the probability of causality (the prediction score) for each SNP, the corresponding Bayes factor (BF_annot_) and scripts to combine BF_annot_ with GWAS association signals.

## Results

### Functional enrichment in GWAS hits

Frequencies of functional characteristics in GWAS hits compared to non-hits were compared using Fisher’s exact test. Our analyses indicate that GWAS hits are enriched for most functional characteristics compared to GWAS non-hits, except for splice sites and micro RNA (miRNA) targets, perhaps due to the very low frequency of these two classes of functional characteristics compared to the others (Table 1 and Table 2).

**Table 1.**
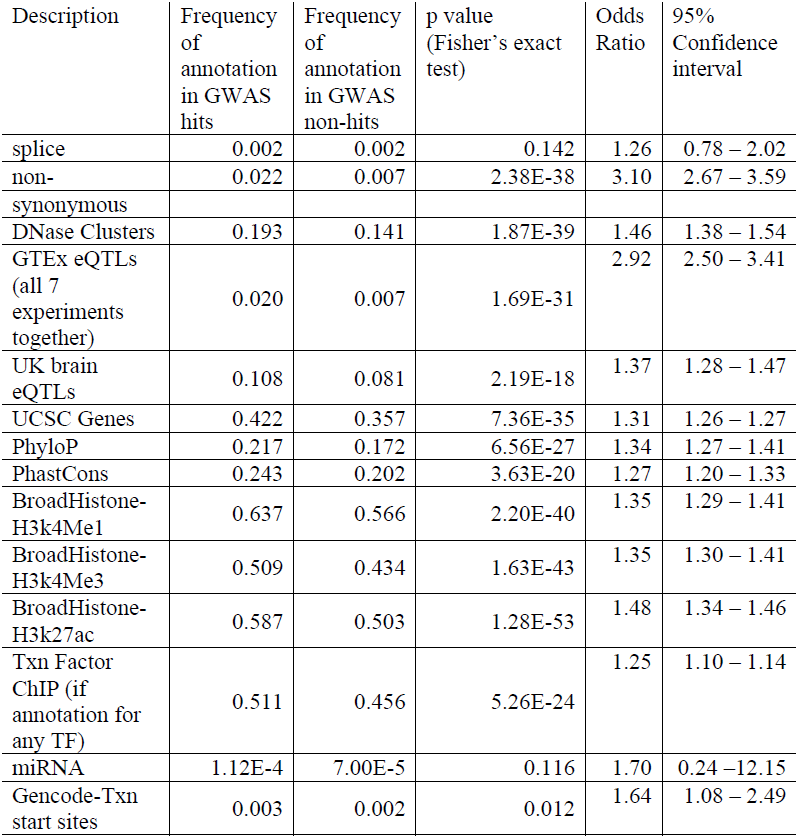
Summary statistics for the functional characteristics in the clumped non-phenotype specific analysis.

**Table 2.**
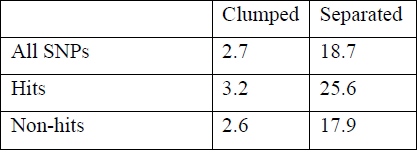
The mean score per SNP across all functional characteristics, classified by SNP type and functional variable type.

|  | Clumped | Separated |
| --- | --- | --- |
| All SNPs | 2.7 | 18.7 |
| Hits | 3.2 | 25.6 |
| Non-hits | 2.6 | 17.9 |

The histone modification data from the Broad Institute had the highest frequencies in GWAS hits, and the lowest p-values for enrichment. Many functional characteristics, most notably miRNA, were very infrequent, but the general picture was that their frequency in GWAS hits was greater than in GWAS non-hits.

We examined the correlations among the various functional characteristics (Figure 1 and Figure 2). The separated-variable analysis included measures of functional characteristics from different cell lines as individual factors, whereas the clumped-variable analysis grouped data from different cell lines for the same functional characteristic. The clumped analysis showed a strong correlation between the two conservation measures (PhyloP and PhastCons), as well as strong correlations among the three histone marks (H3k4Me1, H3k4Me3 and H3k27Ac), and to a lesser degree among the histone marks and transcription factor binding sites. The separated analysis revealed additional correlations among cell types investigated for the DNase I hypersensitive characteristics from Duke University, and to a lesser degree among the DNase I hypersensitive characteristics from the University of Washington, and between these two groups. These results highlight the issue of correlations among functional characteristics, many of which simply represent the same genomic feature, for example a promoter element measured by different technologies. One advantage of elastic net as a regularized logistic regression method is its ability to accommodate highly correlated variables.

**Figure 1.**
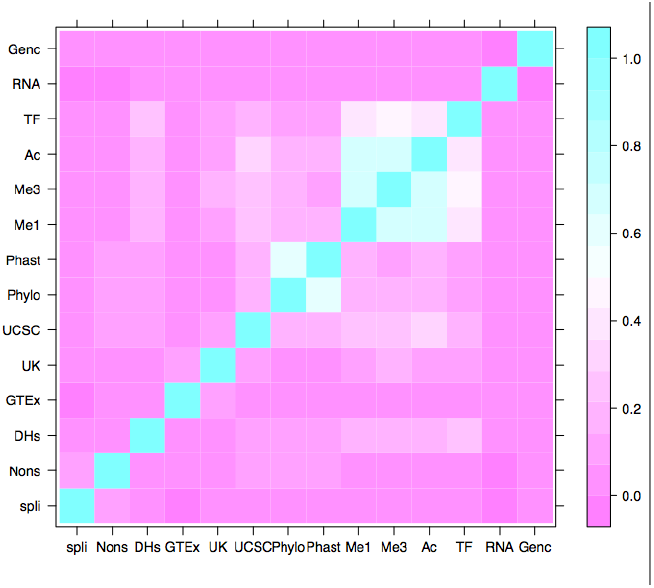
Heat map of correlations among the clumped functional characteristics. High correlations are seen between the two conservation measures PhyloP and PhastCons (represented as Phylo and Phast, respectively). Correlations are also seen among the histone modifications, H3k4Me1, H3k4Me3 and H3k27Ac (Me1, Me3 and Ac, respectively.) Transcription factor binding sites also show a correlation with the histone modifications. [spli = splice sites, Nons = nonsynonymous SNPs, DHs = DNase I hypersensitive sites, GTEx = cis-eQTL data from the GTEx Consortium, UK = cis-eQTL data from the UK Brain Consortium, Phylo = PhyloP conservation, Phast = PhastCons conservation, Me1 = H3K4Me1 histone modification, Me3 = H3K4Me3 histone modification, Ac = H3K27Ac histone modification, TF = transcription factor binding sites, RNA = micro RNA targets, Genc = transcription start sites from Gencode]

**Figure 2.**
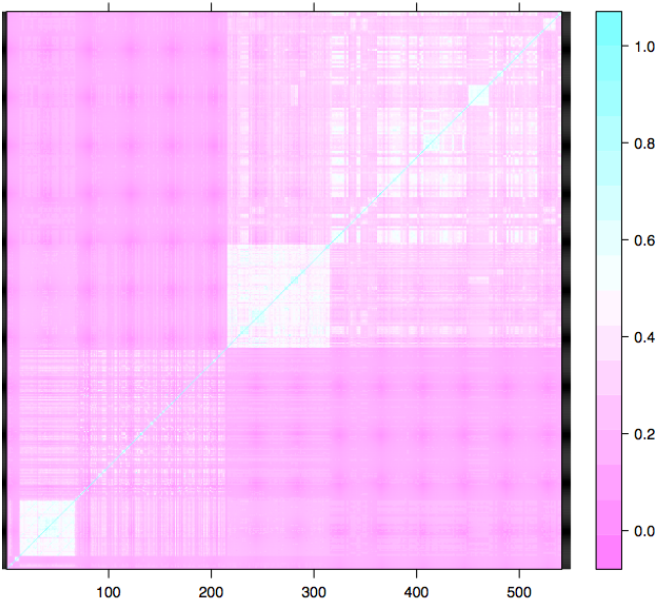
Heat map of correlations among the separated functional characteristics. A full list of the numbered characteristics is provided in Supplementary Table A. The white box in the bottom left corner corresponds to high correlation among the histone modifications. The less defined white area spanning from 72 to 219 on the x axis corresponds to correlation among the transcription factor binding sites, which also show some correlation with the histone modifications. The white box from 220 to 319 on the x axis corresponds to a high correlation among the different cell types for the DNase I hypersensitivity characteristic from Duke University. The less refined white box from around 320 and onwards on the x axis corresponds to the DNase I characteristics from the University of Washington. The plot also shows some correlation among the DNase I characteristics from both groups.

### Predictive accuracy of functional characteristics

We fitted predictive models for GWAS hit status via elastic net, using clumped and separated functional variable sets, using high-confidence (p<5×10^−8^) and low-confidence (p<10^−5^) GWAS hits, and using all GWAS hits (“non-phenotype specific”) as well as hits classified according broad phenotype areas. We primarily investigated predictive accuracy in a separate test set that was not involved in the fitting of the models.

For all of our fitted models, the area under the curve (AUC) of a receiver-operating characteristic (ROC) curve was similar in the test and training sets, suggesting that the models had not been over-fit (data not shown).

We found that the ROC curves for both the separated and clumped analyses had similar AUCs: for instance 0.58 in the test set for the non-phenotype specific clumped analysis and 0.59 in the test set for the separated analysis.

Two analyses emerged as most predictive based on integrating results from ROC curves, positive predictive values, and histograms of the probabilities of causality (the prediction scores). These were the analyses based on non-phenotype specific and the autoimmune GWAS analyses. Best results were obtained from analyses using high-confidence GWAS hits. Results for clumped and separated functional variables were very similar (Table 3 and Figure 3).

**Table 3.** Areas under fitted ROC curves, for clumped-variable analyses. Main values are for analyses of high-confidence GWAS hits. Values in parentheses are for all SNPs in the GWAS Catalogue.

|  | Non-phenotype<br>specific | Brain-related | Cancer | Cardiovascular | Autoimmune |
| --- | --- | --- | --- | --- | --- |
| N | 4480 (8219) | 530 (1741) | 300 (607) | 369 (716) | 570 (863) |
| AUC<br>clumped | 0.68 (0.58) | 0.62 (0.52) | 0.67 (0.60) | 0.69 (0.61) | 0.71 (0.67) |
| AUC<br>separated | 0.70 (0.59) | 0.61 (0.51) | 0.68 (0.60) | 0.66 (0.61) | 0.75 (0.71) |

**Figure 3.**
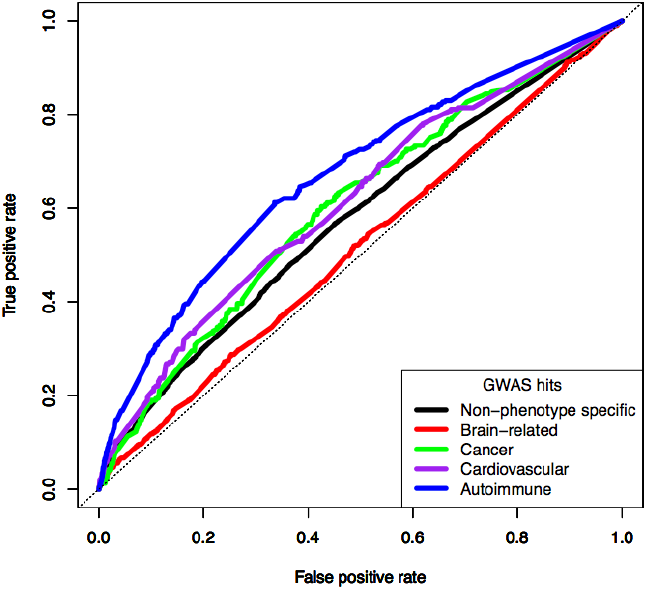
Receiver-operating characteristic (ROC) curves for analyses of clumped functional variables and high-confidence GWAS hits. ROC curves were obtained from a separate test set.

We also investigated positive predictive values (PPVs) and histograms of the probability of causality (prediction score). Overall, the non-phenotype specific analysis using high-confidence GWAS hits produced the highest PPVs as the threshold for declaring a positive hit increased (Figure 4). Histograms of the probability of causality in the test data allowed visualization of the separation (or non-separation) of true hits versus non-hits. We found that for the non-phenotype specific analysis and for the autoimmune analysis, the use of high-confidence GWAS hits in the training data improved the separation of true hits from non-hits in the test data (Figure 5).

**Figure 4.**
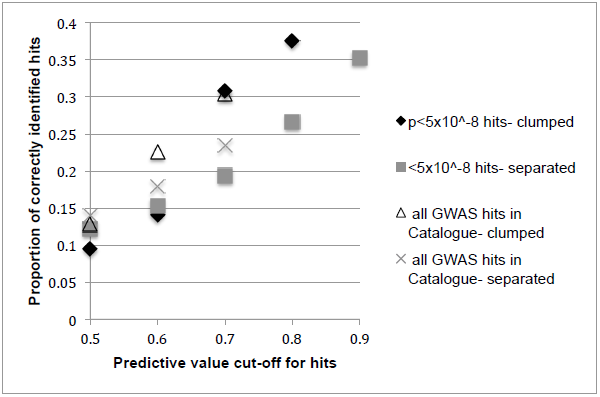
Proportion of correctly identified hits in the test data (positive predictive values). In the non-phenotype specific analyses at various cut-offs for defining hits: SNPs with predictive values of greater than 0.5, 0.6, 0.7, 0.8, or 0.9. Note that results are only plotted for those predictive value thresholds in which there are at least 12 hits correctly identified.

**Figure 5.**
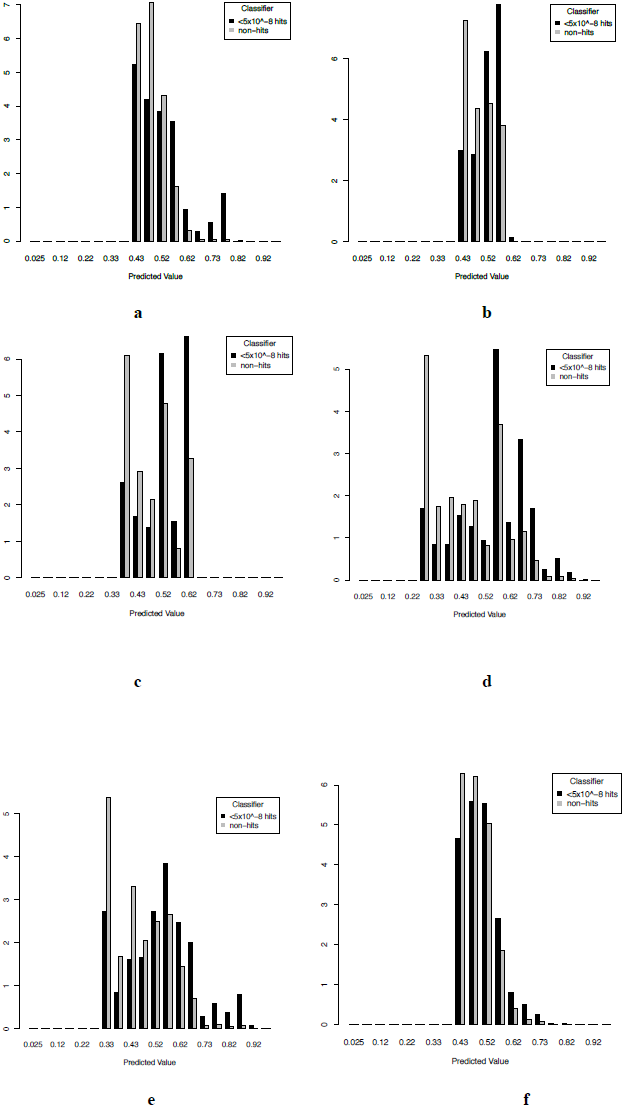
Predicted values for true GWAS hits and non-hits in the test data. Panels show results of clumped-variable analyses on high-confidence GWAS hits for brain-related [**a**], cardiovascular [**b**], cancer [**c**], autoimmune [**d**], and non-phenotype specific hit sets [**e**], and for all hits in the GWAS Catalogue for the non-phenotype specific hit set [**f**].

The results from the histograms of the predicted values from the model were in accord with the PPV results: the non-phenotype specific clumped analysis on high-confidence GWAS hits separated true hits from non-hits most, with the modes of the two distributions clearly distinct. These results suggest that the weighted elastic net procedure was successful in producing models that performed well in identifying true hits as well as in identifying true non-hits. While we could not obtain reliable PPV estimates for the autoimmune analysis due to insufficient data, considering that the PPV results in the non-phenotype specific analysis mirrored the results from the histogram of predicted scores, the separation of non-hits from hits in the histogram was taken as sufficient evidence that the high area under the ROC curve for the autoimmune clumped analysis was also due to positive predictive power.

For the non-phenotype specific clumped analysis, the highest Bayes factor for annotation (22.84) was obtained for rs11177, which is a known GWAS hit associated with osteoarthritis on chromosome 3. It had a predicted value of 0.96. This SNP held all functional characteristics except three low-frequency characteristics: splice sites, miRNA targets, and Gencode transcription start sites. Thirty-five percent of the variants with the top 500 Bayes factors were known GWAS hits. The frequency of hits in the full data was 0.44%. The mean and median of the predicted values for the true hits in the test set were higher than those for the true non-hits (for hits: mean = 0.55 and median = 0.54; for non-hits: mean = 0.46 and median = 0.45).

For the autoimmune clumped analysis, the SNP with the highest Bayes factor was the same as for the non-phenotype specific clumped analysis, rs11177.

### Investigation of the relative importance of different functional characteristics

The importance of a particular functional characteristic in predicting whether or not a SNP is more probable to be a GWAS hit is assessed by means of the magnitude of the coefficient assigned to the characteristic. In both the non-phenotype specific and autoimmune analyses we note that the nonsynonymous SNP functional characteristic had one of the highest coefficients (Figure 6). (The coefficients for both models are provided in **Supplementary Information Part A**.) GTEx eQTLs had the highest coefficient in the autoimmune analysis.

**Figure 6.**
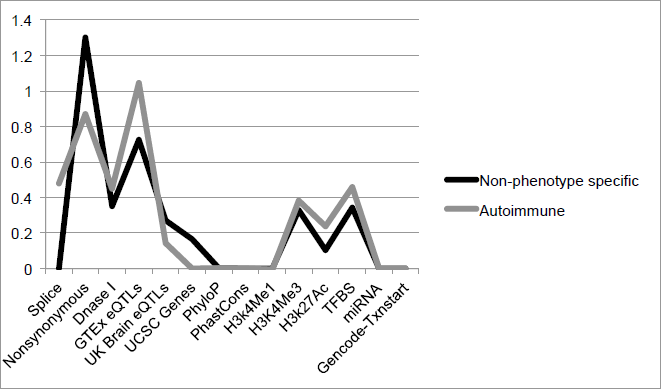
Coefficients of the functional characteristics for the two best analyses. The figure shows the coefficients from the clumped analysis on high-confidence GWAS hits for the non-phenotype specific versus the autoimmune model.

### Investigating functional predictions in the context of known GWAS

We investigated: schizophrenia from a meta-analysis GWAS involving the first sample from the Psychiatric Genomics Consortium (PGC1) combined with a Swedish sample [15], systolic blood pressure from the International Consortium for Blood Pressure (ICBP) [16], and height from Genetic Investigation of Anthropomorphic Traits (GIANT) Consortium [17]. The studies analyzed over 35,000 cases and 47,000 controls, 200,000 individuals and, and over 180,000 individuals, respectively.

For each study, we stratified the quantile-quantile plots according to predicted value bins (Figure 7). We found that SNPs with higher predicted values from the non-phenotype specific clumped analysis tended to deviate more from the line corresponding to the overall GWAS, in favour of more association signals. Similar results were obtained for all three GWAS analyzed: schizophrenia, systolic blood pressure and height.

**Figure 7.**
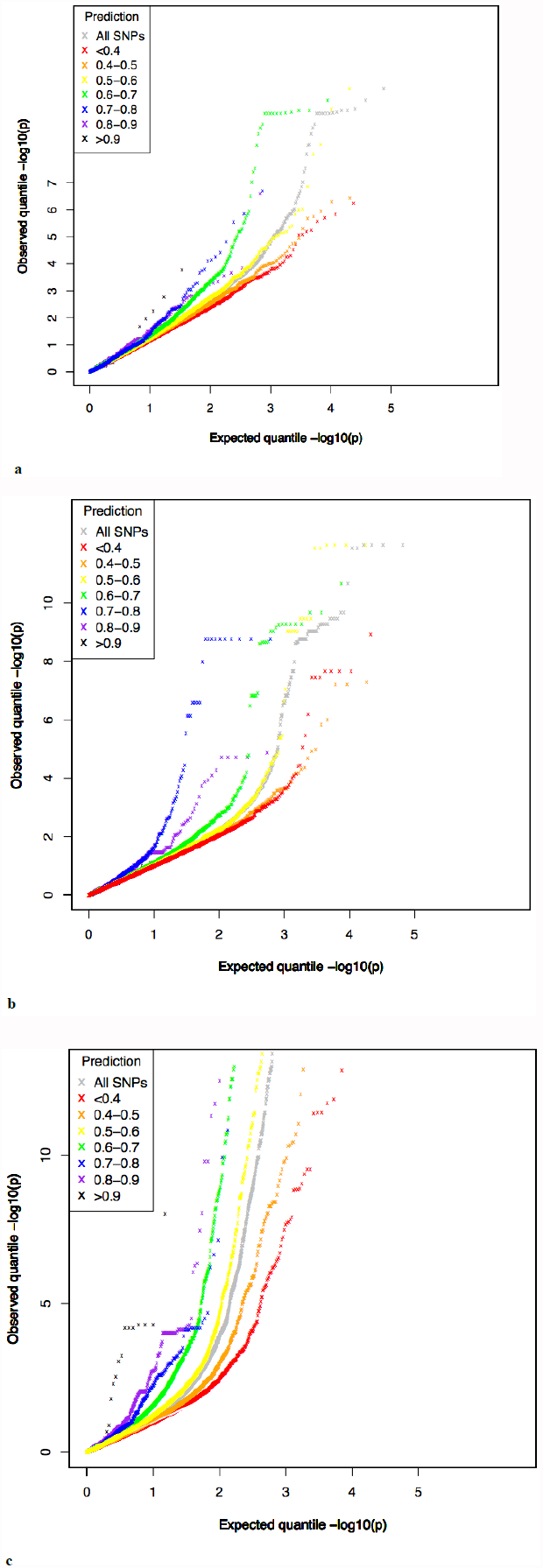
Quantile-quantile plots stratified by predicted values for SNPs in real GWAS. All GWAS SNPs (in grey) for a schizophrenia GWAS from PGC1 with a Swedish sample [**a**], a systolic blood pressure GWAS from ICBP [**b**], and a height GWAS from GIANT [**c**]. The non-grey lines show plots for SNPs binned according to their predicted value from the non-phenotype specific model.

We obtained summary data obtained from a psoriasis GWAS study from Strange et al. [18]. We then selected 15 SNPs that were subsequently discovered in a meta-analysis [19]. Using summary association statistics from the Strange et al. study we derived Bayes factors for association (BF_assoc_) and Bayes factors based on association data combined with the annotation of functional characteristics (BF_assoc_*BF_annot_) for each SNP. We ranked the SNPs according to BF_assoc_, and ranked them again according to BF_assoc_*BF_annot_ to determine whether annotating SNPs with their functional characteristics improved their rank (larger Bayes factors were assigned smaller ranks). BF_annot_ values were derived from the non-phenotype specific clumped analysis using high-confidence GWAS hits. As negative controls, we took 8 independent sets of a random 15 SNPs (which were not in high LD with any of the 15 hits and had similar p-values to the hits) and compared the difference in the sum of ranks based on BF_assoc_ versus BF_assoc_*BF_annot_. The procedure was repeated using BF_annot_ derived from the autoimmune clumped analysis.

Of the 15 true psoriasis hit SNPs, 7 had better ranks based on BF_assoc_*BF_annot_ compared to association information on its own (BF_assoc_). The difference of the sum of ranks assigned to the 15 hits was nearly 48,000 based on BF_assoc_*BF_annot_ compared to BF_assoc_, with the former having the lower sum (better ranks). Many of the hit SNPs had very large ranks based merely on the association data (>3000), which was also the case for ranks based on BF_assoc_*BF_annot_, but the trend was in the right direction with better ranks obtained when combing the association information with the annotation of functional characteristics. Of the 12 random sets of 15 independent SNPs, the trend was in the opposite direction for 10 of the sets (with SNPs having better ranks based on BF_assoc_ alone). Of the remaining 2 sets, one of them had the same number of the SNPs with improved ranks based on BF_assoc_*BF_annot_ compared to BF_assoc_ as did the analysis with the actual hits (7 out of 15), and the other random set had 8 SNPs that showed improvement. However, for those random SNP lists the difference in the sum of ranks from BF_assoc_ compared to BF_assoc_*BF_annot_ was less than half of the improvement of ranks seen for the 15 hits. Comparable results were seen when using BF_assoc_ based on the autoimmune clumped analysis. The difference between the sum of the ranks for BF_annot_ compared to BF_assoc_*BF_annot_ was over 49,000, with improved ranks of the hits based on the BF_assoc_*BF_annot_ ranks. Of the random lists the largest difference in the sum of ranks from BF_assoc_ compared to BF_assoc_*BF_annot_ was less than a third of the improvement of ranks seen for the 15 hits.

## Discussion

The release of major genome wide datasets such as ENCODE and NIH Roadmap projects, offers a unique opportunity to re-assess the existing GWAS corpus and draw conclusions about which functional characteristics in the human genome are most likely to indicate causality in association studies. We previously considered Bayes factors based on a limited set of functional characteristics, considering each functional characteristic separately [3]. Here we have extended our Bayesian framework by developing Bayes factors for multiple functional characteristics, considering all functional characteristics jointly. We used a regularized logistic regression to fit predictive models allowing for large numbers of both qualitative and quantitative functional characteristic data. We performed our analysis under a wide variety of conditions, including phenotype specific analysis for autoimmune, brain-related, cancer, and cardiovascular disorders.

Our results confirm previous findings of differences in functional enrichment in GWAS hits compared to non-hits, which provided a rationale for utilizing functional characteristics as predictors of SNP causality. We found that using high-confidence GWAS hits (p<5×10^−8^) as a classifier resulted in more predictive power. However, if the number of GWAS hits that are available for training are too low, then the predictions become imprecise. This was a reoccurring theme for many of the phenotype specific analyses. The separation between true GWAS hits and non-hits in the test set, in addition to the AUC, should be used to assess the predictive power of a model. Using those methods we found that the non-phenotype specific and the autoimmune analyses on clumped variables using high-confidence GWAS hits were most reliable. For instance, although the AUCs were slightly higher for the separated analyses, the classification of true GWAS hits and non-hits was better in the clumped analysis, suggesting that the clumped analysis may provide more accurate predictions. The benefit of the separated analysis is that it allows researchers to identify characteristics specific to certain conditions, for example specific cell types, which can be useful for planning further investigations, but the increased number of variables and sparsity of the data reduces the power of this type of analysis.

A limitation to the study is the restricted amount of tissue- or cell-specific data, especially in light of the findings that enrichment of disease-specific GWAS hits can differ in certain cell types, for example for DNase I hypersensitive sites [8]. Incorporating additional functional characteristics, for example those from relevant tissue types, will likely improve the understanding of which characteristics are associated with GWAS hit SNPs, especially for the phenotype specific analyses.

The current number of GWAS hits in the GWAS Catalogue makes it challenging to sub-divide hits into phenotype specific traits. However, preliminary results showing differences in the coefficients for the functional characteristics suggest that as the number of GWAS hits grows, a phenotype specific approach from which to derive Bayes factors for prioritization could be more biologically relevant than simply an approach that combines all GWAS hits together. Interestingly, although it was one of the largest lists, the brain-related list did not have a greater predictive power than expected by chance. This finding only serves to reinforce the widely appreciated complexity of brain-related disorders. Nevertheless, schizophrenia GWAS significant SNPs showed enrichment of SNPs with high predicted values from the model, as did SNPs associated with systolic blood pressure or height.

Using manually curated phenotype lists as done here may not be the best option. Using lists that are more reproducible, such as those based on the Experimental Factor Ontology (EFO) definitions, may be more appealing. However, most of the lists created using the EFO definitions were relatively small, covering less than 10% of the total GWAS hits on the common genotyping arrays, and thus this method of classifying GWAS hits was deemed to be not feasible, but may be possible in the future as the size of GWAS Catalogue grows still larger.

The coefficient for non-synonymous SNPs was the highest in the non-phenotype specific analysis and a close second in the autoimmune analysis. This result suggests that being a variant in a gene that causes a protein alteration is an important indicator of whether or not a genetic variant will be truly associated with a phenotype. The result agrees with the findings that the top associated SNPs and also those that are nominally associated with a phenotype are more likely to overlap genes than non-GWAS SNPs [20]. GTEx eQTLs came up as the most important factor in the autoimmune analysis. Two of the experiments analyzed eQTLs from lymphoblastoid cells, which may explain the importance of this functional characteristic in the autoimmune traits.

We have shown that our method can be used to calculate Bayes factors for annotation (BF_annot_). These can be applied to GWAS data to prioritise near-significant variants for follow-up based on the likelihood of being causal in light of their functional characteristics. The method takes LD into account, and uses information from the March 2012 release of the 1000 Genomes Project to map relevant annotation information from all variants in high LD, including both SNPs and indels. In addition to being used for variant prioritization of GWAS data, the methodology could be applied in the future to the prioritization of variants from fine mapping and sequencing studies. Here, the question arises as to whether the models described here, which were created based on common variation, could be applied to rare variation. In time, larger databases of true causal variation, including rare variation, will allow our method to be applied with increasing accuracy.

## Methods

### Representative GWAS SNPs

To represent the characteristics of a typical GWAS panel, markers from the Affymetrix Genome-Wide Human SNP Array 6.0, the Illumina Human1M-Duo Genotyping BeadChip, and the Illumina HumanOmni1-Quad BeadChip were downloaded from the UCSC genome browser, using the table browser tool [21]. The union of these three arrays consisted of 1,936,864 unique SNPs from the 22 autosomes. Because of its unique LD and genic properties, the MHC region (chr6:29624809 - 33160245 on build 37) was excluded from downstream analyses.

LD proxies or “tagging” SNPs (r^2^>=0.8) for the GWAS panel SNPs were identified using VCFtools [22] based on data from the (N=379) Europeans (Phase I, version 3, March 14, 2012) in the 1000 Genomes Project [23].

GWAS “non-hits” were defined as all those SNPs in our union GWAS set which were neither a GWAS “hit” (see below), nor in high LD (r^2^>=0.8) with a GWAS hit.

### GWAS hits

To obtain a set of SNPs (and their LD proxies) with good prior evidence of causality, we downloaded the Catalogue of Published Genome-wide Association Studies from the National Human Genome Research Institute (NHGRI) (http://www.genome.gov/gwastudies) [1] on August 6, 2013. This catalogue contains a list of SNPs that have been shown to be associated with a particular trait in a GWAS at a suggestive p-value <10^−5^. We removed SNPs in the Catalogue that were not present in the representative GWAS set defined above, and similarly removed SNPs on the sex chromosomes or in the MHC region.

### Functional characteristics

We acquired functional data from a variety of sources (Table 4). A full list is provided in **Supplementary Table A**. Much of the data was downloaded from the UCSC genome browser using the table browser tool [21]. Additionally, a substantial proportion of the data was derived from the ENcyclopedia of DNA Elements (ENCODE) Project Consortium, which developed and implemented a range of experimental techniques with the aim of identifying the functional regions of the human genome, particularly including non-coding regions [24]. Data from this project that were used included transcription factor binding sites (TFBSs), three histone modifications (H3K4Me1, H3K4Me3, H3K27Ac), and DNase I hypersensitive sites. H3K4Me1 is associated with enhancers and DNA regions downstream of transcription starts, and often found near regulatory elements; H3K4Me3 is associated with promoters active or poised to be active, and often found near promoters; H3K27Ac thought to enhance transcription possibly by blocking repressive histone mark H3K27Me3, and often found near active regulatory elements. DNase I hypersensitive sites are regions in the genome with high affinity of being cleaved by the DNase I enzyme. The technologies for identifying the functional characteristics mentioned above used chromatin immunoprecipitation followed by sequencing (ChIP-seq), with the exception of the University of Washington (UW) group that identified DNase I hypersensitive sites using Digital DNase I. This method involves DNase I digestion of intact nuclei, isolation of DNaseI “double-hit” fragments, and direct sequencing of fragment ends. Peaks are regions that are enriched in the captured fraction of the DNA suggesting they are occupied by the protein of interest (any score > 0). We used a binary variable to indicate whether a SNP was within a peak.

**Table 4.** Summary of functional characteristics.

| Functional characteristic analysed | Description | Number and detail of measures used in the analysis* |  |
| --- | --- | --- | --- |
|  |  | Clumped | Separated |
| ENCODE data |  |  |  |
| UW DNase I hypersensitive sites | Data from digital DNaseI methodology; (“peaks”) | N/A | 226 |
| Duke DNase I hypersensitive sites | Positions of open chromatin by Formaldehyde-Assisted Isolation of Regulatory Elements (FAIRE) and ChIP-seq experiments; (“peaks”) | N/A | 100 |
| DNase Clusters (v2)** | Stringent (FDR 1% threshold) for “peaks” of DNase I hypersensitivity from uniform processing by the ENCODE Analysis Working Group of data from UW and Duke | 1 | N/A |
| Txn Factor ChIP | Transcription factor binding sites (TFBS) from ChIP Seq experiments; (“peaks”) | 1 (presence or absence in any TFBS) | 148 (separated by TF, but <u>not</u> by cell type due to sparse data) |
| Broad Histone – H3K4Me1, H3K4Me3, H3K27Ac | All are assayed using ChIP-Seq; (“peaks”) | 3 (each histone mark grouped by the 18 cell types and/or conditions) | 54 (each histone mark separated by cell type and/or conditions) |
| Conservation |  |  |  |
| PhyloP | Average scores can be calculated as the sum of scores divided by the number of valid data values in the block (scores range from 0.1 to 2.2910) | 1 | 1 |
| PhastCons | Average scores can be calculated as for PhyloP (scores range from 0.1 to 1.0 in this dataset) | 1 | 1 |
| Expression quantitative trait loci |  |  |  |
| eQTL- GTEx | cis-eQTLs, $p<1\times10^{-5}$ cut-off for variants within 2Mb of the expressed gene. | 1 (any eQTL) | 7 (separated by dataset) |
| eQTLs - UK Brain | cis-eQTLs, FDR<1% cut-off for variants within 2Mb of the expressed gene. | 1 | 1 |
| Other characteristics |  |  |  |
| UCSC Genes | UCSC known Gene | 1 | 1 |
| Splice sites | Splice site region defined as -5 to +5 range around exon starts & exon ends of UCSC Genes | 1 | 1 |
| Nonsynonymous SNPs | Coding Nonsynonymous SNPs defined as stop-gain (nonsense), missense, stop-lost, frameshift or inframe indel | 1 | 1 |
| TS miRNA sites | Conserved mammalian microRNA regulatory target sites for conserved microRNA families | 1 | 1 |
| Gencode | Based on the GENCODE Genes variable | 1 | 1 |
| transcription start sites | (version 17, June 2013) |  |  |

\* All SNPs are annotated in a binary fashion indicating the presence or absence of a functional characteristic, except for the conservation scores, for which the SNPs are assigned a quantitative score.
\*\* The DNase Clusters v2 file was created by combining the UW and Duke DNase I data that have been uniformly processed and replicates merged. Stringent (FDR 1% thresholded) peaks of DNase I hypersensitivity from uniform processing by the ENCODE Analysis Working Group were applied. Grouping the UW and the Duke DNase I hypersensitive variables are not equivalent to the DNase Clusters v2 file, and thus we used the latter to represent DNase I hypersensitive sites in the clumped analysis due to the substantial efforts made to combine the data meaningfully.

Two types of conservation scores from 46 placental mammals (PhyloP and PhastCons) were incorporated. Both PhyloP and PhastCons scores are derived using phylogenetic hidden Markov models. These two measures have their own advantages. PhyloP scores do not take into account conservation at neighbouring sites, whereas PhastCons estimates the probability that each nucleotide belongs to a conserved element.

Expression quantitative trait loci (eQTLs), which are variants that are correlated with gene expression, were included. In particular those that fall within 2Mb (+/-1Mb upstream and downstream) (cis-eQTLs) of the gene of interest were used. These data were derived from the NCBI-hosted GTEx Browser (http://www.ncbi.nlm.nih.gov/gtex/GTEX2/gtex.cgi) and the UK Brain Expression Consortium [25].

Summary information concerning the location or function within a gene (coding-non-synonymous, coding-synonymous, splice site, untranslated regions, etc) was derived from dbSNP. Non-synonymous SNPs, were classified as those SNPs with one of the following characteristics: stop-gain (nonsense), missense, stop-lost, frameshift or inframe indel. Splice site regions were defined as being within five base pairs upstream and five base pairs downstream of the exon start site or the exon end site. The UCSC gene table was used to determine the exon start and end sites. The UCSC gene table is comprised of a set of gene predictions based on data from RefSeq, GenBank, the Consensus Coding Sequence (CCDS) variable, Rfam, and the Transfer RNA Genes variable. Additional characteristics used were 3’ targets for microRNA (miRNA), and also transcription start sites as described by Gencode [26]. As miRNA targets are known to be substantially over-predicted, we used a conservative miRNA target dataset based on conserved mammalian microRNA regulatory target sites in the 3’ UTR regions of Refseq Genes, as predicted by the TargetScan algorithm (Human 5.1) [27].

All SNPs in our GWAS hit and GWAS non-hit sets, along with all their LD proxies, were annotated with all the functional characteristics defined above. Each GWAS hit and non-hit SNP was then given the maximum value for each functional characteristic found across all its LD proxies.

### Tests for functional enrichment

Counts of GWAS hits and non-hits were categorized by annotation value and compared using Fisher’s exact test. To verify that results were not unduly influenced by correlations (LD) among observations, we also conducted analyses in which genetic variants were “pruned” so that all SNPs have r^2^<0.8 with all other SNPs. The results of these analyses were very similar (data not shown).

Heat maps were constructed using R [28] to compare correlations among the various functional characteristics.

### Regularized logistic regression via elastic net

We used a regularized form of logistic regression known as elastic net to predict GWAS hit versus non-hit status on the basis of the functional characteristics we had collected. We first employed this method for a symposium on “Functional annotation of GWAS hits” that we organized for the American Society of Human Genetics in 2010. Elastic net is a form of machine learning first described by Zou and Hastie [29], and is implemented in the glmnet package [20] in R [28]. Briefly, regularization is achieved via the subtraction of a penalty term from the log-likelihood prior to maximization. The penalty term includes both a “lasso-like” L1 component (the sum of the absolute values of all fitted coefficients) and a “ridge-like” L2 component (the sum of squares of all fitted coefficients). Two parameters, alpha and lambda, determine the relative importance of the L1 versus the L2 term (alpha), and the overall importance of the penalty term in the maximization (lambda). Appropriate values for these parameters were found by 10-fold cross-validation of the training set (see below).

Due to the unbalanced nature of the data (many more GWAS non-hits than hits) we employed a weighting procedure in the logistic regression to balance the accuracy of prediction in both types of markers. We weighted all hits by (N_hits_+N_non-hits_)/2N_hits_ and all non-hits by (N_hits_+N_non-hits_)/2N_non-hits_, where N_hits_ and N_non-hits_ denote the number of hits and non-hits, respectively, in the training set. This procedure has the effect of equalizing the importance of hits and non-hits in the logistic regression.

We randomly selected 60% of our GWAS hits and non-hits to form our training set. The remaining 40% of the data (the test set) was used to assess the performance of the model using ROC curves and other measures. We repeated the machine learning modifying the percentage of the data used in the training and test sets, and all splits produced similar results (**Supplementary Information Part B**). The 60%/40% training/test set split was pursued for the remaining analyses.

The data was split into the training and test sets ten times using a random number generator. We found that the beta coefficients were consistent for all of the functional characteristics with the exception of those with the lowest frequencies (**Supplementary Information Part C**).

For the calculation of Bayes factors, we performed elastic net, using the same determined values of alpha and lambda, on the full GWAS hit and non-hit datasets.

### Predictive accuracy

We employed three methods to determine which models had the best predictive accuracy: ROC curves, positive predictive values, and histograms of the predicted values from the models.

ROC curves show the sensitivity and specificity of a fitted model. Sensitivity is the probability of the model providing a true positive result (identifying a true GWAS hit in the test set). Specificity is the probability of the model providing a true negative result (identifying a true GWAS non-hit in the test set). An AUC of 0.5 indicates a model of no predictive value, while an AUC of 1 indicates perfect predictive power. The ROC curves were created using the ROCR package [31] in R.

ROC curves do not reflect how well a model performs within each class given unbalanced data (a very large number of non-hit SNPs compared to hits). To capture this aspect we also investigated positive predictive values (PPVs), the proportion of SNPs with predicted probabilities of causality above a certain threshold (we investigated thresholds of 0.5, 0.6, 0.7, 0.8 or 0.9) that are true GWAS hits in the test set. Finally, we visualized class separation with histograms of the predicted probabilities of causality by class.

### Definition of functional variables and GWAS hits

A variety of functional characteristics were investigated as input variables. One, defined as the “clumped” analysis, featured groups of functional characteristics, which were collapsed into a single summary variable. The “separated” analysis worked on all functional characteristics individually.

We performed phenotype specific analyses in which the analyses outlined above were carried out using phenotype specific GWAS hits as classifiers. An autoimmune list, a brain-related list and a cardiovascular list were created using the GWAS Catalogue searching for terms relating to those phenotypes. Each list was then verified by an expert in the field.

Additionally, the GWAS Catalogue was divided up into categories specified by the Experimental Factor Ontology (EFO) definitions; however, due to small numbers of SNPs in each category this mode of classification is not currently feasible for most of the subsets (**Supplementary Information Part D**). Only the cancer list, which was the largest disease-relevant list, was used.

We defined two sets of GWAS hits for downstream analysis, one based on a weak significance threshold of p<10^−5^ and one based on a strong significance threshold of p<5×10^−8^, as reported in the NHGRI GWAS Catalogue.

### Derivation of Bayes Factors

Bayesian analysis provides the most suitable framework for combining functional characteristics (here referred to as “annotation data”), with evidence from an association study (“association data”) [32]. We expand on our previous empirically-based approach to the calculation of Bayes factors for annotation [3] to allow multiple functional characteristics to be considered simultaneously. The posterior odds (O_post_) of causality for a trait of interest at a given SNP are given by the ratio of the conditional probability of causality, given the annotation and the association data, to the conditional probability of non-causality:

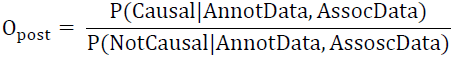

If we assume the annotation data and association data are independent once conditioned on causality, then the posterior odds become:

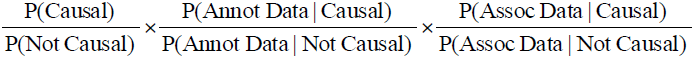

These three products are, respectively, the prior odds before seeing any association and annotation data (O_prior_), the Bayes factor for annotation data (BF_annot_) and the Bayes factor for association data (BF_assoc_). We note that this factorization implies that, while functional annotations are allowed to be enriched (or impoverished) for causal SNPs relative to non-causal SNPs, the enrichment pattern is assumed to be the same for rare versus common causal SNPs, and for low-effect size versus high effect size causal SNPs. We accept that this is an imperfect approximation, and it assumes among other things that SNPs are either causal or non-causal when in reality their effect size can be arbitrarily close to zero, but we note that the main limitation of our approach lies with the small number of GWAS hits available to us, and subdividing these still further according to allele frequency and effect size would be problematic. We also note that by “causal” what we actually mean is “causal or in high LD with a causal variant”, as both the association data and the annotation data (as defined in our study) are affected by LD proxies.

In our previous study [3], we noted that if one assumed that (1) all hits in the NHGRI GWAS Catalogue were truly causal; and (2) functional annotation enrichment patterns were the same for these known hits as for future undiscovered truly causal SNPs; then an empirically based estimate for BF_annot_ for a single binary functional characteristic would simply be the ratio of its frequency in GWAS hit versus non-hit data. Here we note that if we start with the same two assumptions, and further assume that a true (but unknown) logistic model exists that relates a set of functional characteristics (which can be either binary or quantitative) to the probability that a SNP is truly causal, then one reasonable approach to estimating that logistic model would be via regularized logistic regression as described above. Once fitted, the estimated odds of causality to non-causality, obtained from the GWAS hit and non-hit datasets, need only be multiplied by the prior odds of non-causality in these dataset (i.e. the ratio of the weighted sample sizes of GWAS non-hits to GWAS hits in these data) in order to obtain the Bayes factor for annotation. Here, we chose to weight hits and non-hits to appear of equal size, and thus our estimate for BF_annot_ is obtained directly as the estimated odds of causality to non-causality from the regularized logistic regression.

Methods for estimating BF_assoc_ from association data are reviewed by Stephens & Balding [32]. Here, we use the convenient approximation described by Wakefield [33].

### Investigating the model in the context of known GWAS

To investigate the relevance of the predictions in a variety of disorders we looked at the p-value distribution of SNPs according to their functional class in large GWAS datasets with a substantial fraction of GWAS significant findings. Quantile-quantile plots were constructed for each study with multiple lines corresponding to SNPs binned according to their predicted value. Predicted values were those derived from the non-phenotype specific clumped model in which GWAS hits were defined as those SNPs in the GWAS Catalogue with p-values of less than 5×10^−8^. We expected those SNPs with higher predicted values to be enriched with GWAS SNPs with more significant p-values, whereas those SNPs with lower predicted values would be enriched with less significant p-values compared to all SNPs in the GWAS.

We also selected some SNPs shown to be associated in a large psoriasis meta-analysis which had not been identified in a previous GWAS study [18, 19]. We then determined the effect on the rank of their Bayes Factors in the previous study derived either using association data or both association data and functional characteristics.

## Acknowledgements

We would like to thank David Kevans and Mira Ryten for their help in defining the phenotype specific lists. We would also like to thank Andrew Patterson, James Kennedy and Arturas Petronis for comments on the overall project. This work forms part of the research themes contributing to the translational research portfolio of the National Institute for Health Cardiovascular Biomedical Research Unit at Barts, Michael Barnes is funded by this award.

